# Potential mammalian species for investigating the past connections between Amazonia and the Atlantic Forest

**DOI:** 10.1101/2020.12.24.424335

**Authors:** Arielli Fabrício Machado, Camila Duarte Ritter, Cleuton Lima Miranda, Maria João Ramos Pereira, Leandro Duarte

**Author notes:** Author for correspondence: Arielli F. Machado. Postal address: Phylogenetic and Functional Ecology Lab (LEFF), Post-Graduation Programme in Ecology, Universidade Federal do Rio Grande do Sul (UFRGS), Bento Gonçalves Avenue, 9500, Porto Alegre, RS, 91501-970, Brazil. Camila D. Ritter. Postal address: University of Duisburg-Essen, Universitätsstrasse 5 - D-45141 Essen, Germany.

## Abstract

Much evidence suggests that Amazonia and the Atlantic Forest were connected through at least three dispersion routes in the past: the northeast route, the central route, and the southeast-northwest route. According to previous studies, the southeast-northwest route would have been the most frequently used. However, few studies have assessed the use of these routes based on multiple species. Here we present a compilation of potential mammal species that could have dispersed between the two forest regions to investigate these connections. We evaluate the geographic distributions of mammals occurring in both Amazonia and the Atlantic Forest and the likely connective routes between these forests. We classified the species per habitat occupancy (strict forest specialists, species that prefer forest, or generalists) and compiled the genetic data available for each species to evaluate their potential for phylogeographic studies focusing on genetic exchange between the two forest regions. We found 127 mammalian species occurring in both Amazonia and the Atlantic Forest for which significant genetic data was available. Hence, highlighting their potential for phylogeographic studies investigating the past connections between the two forests. Differently from what was previously proposed, the northeast route showed evidence of past use by more mammal species than the remaining two routes. The central route would have been the second most important in terms of species. Our results show the potential of using mammal species to investigate and bring new insights about the past connections between Amazonia and the Atlantic Forest.

## Introduction

Amazonia and the Atlantic Forest are among the most diverse forests in the world [1, 2]. Biogeographical patterns of these megadiverse forests have been investigated since the 19^th^ century [3, 4]. Currently, these forests are separated by the ‘dry diagonal’ comprising the Caatinga, the Cerrado and the Dry Chaco ecoregions. However, different evidence sources including biogeographical [5–10], palynological [11–13], and geological [14] show that these forests have been connected in the past.

Three routes have been suggested as past connections between Amazonia and the Atlantic Forest, one through the forests of North-eastern Brazil (the northeast route [NE route]), another through the gallery forest of the Brazilian Cerrado ecoregion (the central route [CE route]) and a third through the forests of the Paraná Basin, the Moist Chaco, and the Pantanal (the southeast-northwest route [SW route]) [5, 6, 15]. According to Por [5], the SW route would have been the first to be formed and this connection would have occurred more often over time, followed by the NE route.

In a study aimed to test Por’s hypothesis [5], Ledo & Coli [10] reviewed the literature for molecular evidence of connections between Amazonia and the Atlantic Forest for ca. 60 vertebrates, including 10 mammals. They found more studies that evidenced connections through the SW route compared to the NE route [10]. However, this result could be biased due to the poor sampling in the northeast region [16]. It remains uncertain whether the SW route was indeed the most frequently formed connection and thus the most used route in the past.

There is a great number of recent sister species of birds in Amazonia and the Atlantic Forest that may have resulted from the use of the NE route [8]. However, like Ledo & Colli [10], these authors did not consider the CE route as independent but included it as part of the NE route. Yet, the CE route has been well documented as a separate migratory pathway in the literature for both animals and plants [5, 6, 15, 17, 18]. Moreover, investigating eight small mammals, Costa [6] found a larger number of small, related mammals occurring in Amazonia and the Atlantic Forest that could have come through the CE route. However, this study was limited for small mammals which have specific traits, such as limited dispersion ability. Considering more species with different traits and divergence time may further add evidence on the use of the CE connection, as well as for the other routes. In this context, ecological and genetic data may shed light on the use and frequency of these past connections.

Some phylogeographical studies investigate the role of the historical connections between Amazonia and the Atlantic Forest in terms of dispersion and diversification of several animal species, such as mammals [6, 19], birds [8], reptiles [20–22], amphibians [23] (for a literature revision of vertebrate evidence see [10]), and insects [24]. However, the totality of species that may evidence past connections between Amazonia and the Atlantic Forest have not been mapped and this information is particularly scarce for mammals.

There are currently several databases of species geographic and molecular distribution data that can be used to evaluate those past forest connections. A compilation of the data available, including geographic, ecological, and genetic data, for species that may show evidence of the past connections between Amazonia and the Atlantic Forest could thus be especially useful for testing the aforementioned hypotheses.

Here we aim to identify potential mammal species useful for investigating the past connections between Amazonia and the Atlantic Forest through geographical distribution patterns, habitat preference, and genetic data. Furthermore, we aim to identify the potential connective routes previously proposed in the literature. We believe that our results may serve as basis for future biogeographic studies considering different mammalian taxonomic groups.

## Material and Methods

### Species data

We considered mammal species of interest for investigating the past connections between Amazonia and the Atlantic Forest to fulfil the following three criteria: 1) species occurring in both Amazonia and the Atlantic Forest, 2) species that use forests, and 3) species with genetic data available on GenBank [25]. Additionally, we identified potential past connective routes between Amazonia and the Atlantic Forest by investigating the distribution maps for each of these species.

### Geographical data

Geographical distribution maps of forest mammalian species from Amazonia and the Atlantic Forest were compiled from the IUCN - International Union for Conservation of Nature [26]. To identify the mammalian species that occur the two regions, Amazonia and Atlantic Forest, the IUCN maps were overlaid on the Ecoregion maps [27] using the Amazonian and the Atlantic Forest limits, through the *gIntersection* function of the R package ‘rgeos’ v. 0.5.5 [28] in R v. 3.6.3 [29]. Subsequently, only species with occurrences in both Amazonia and the Atlantic Forest were selected. The predefined identifications based on the overlaid IUCN occurrence maps were revised using the annotated list of mammals in Brazil of Paglia et al. [30] since this reference agree with current geographical and genetic data available in the databases used in this study.

### Habitat classification

We selected solely forest species by accessing the IUCN information on species habitat use through the *rl_habitats* function of the R package ‘rredlist’ v. 0.6.0 [31]. Species that were exclusive to open and/or aquatic habitats were not considered for this study. We generated a scale of habitat preference for each species from forest specialist to generalist, as this is key information for studies about the connective routes between Amazonia and the Atlantic Forest. We based this scale of habitat preference on the detailed text about species’ habitat and ecology, available in the IUCN database [25] and additional literature reviews [e.g., 33–38]. The criteria used for classifying the species according to habitat were as follows: 1) Strict forest specialists (SF), encompassing species that only occur in forests, 2) Species that prefer forest (PF), encompassing species that use not only forested habitats but prefer these environments, or 3) Generalists (G) encompassing species that use both forests and open environments (Table 1). Then, we used a Pearson’s Chi-squared test to assess the relationship between the species’ habitat preferences and the routes used through the *chisq.test* function in the R v. 3.6.3 basic package ‘stats’ [28].

**Table 1.**
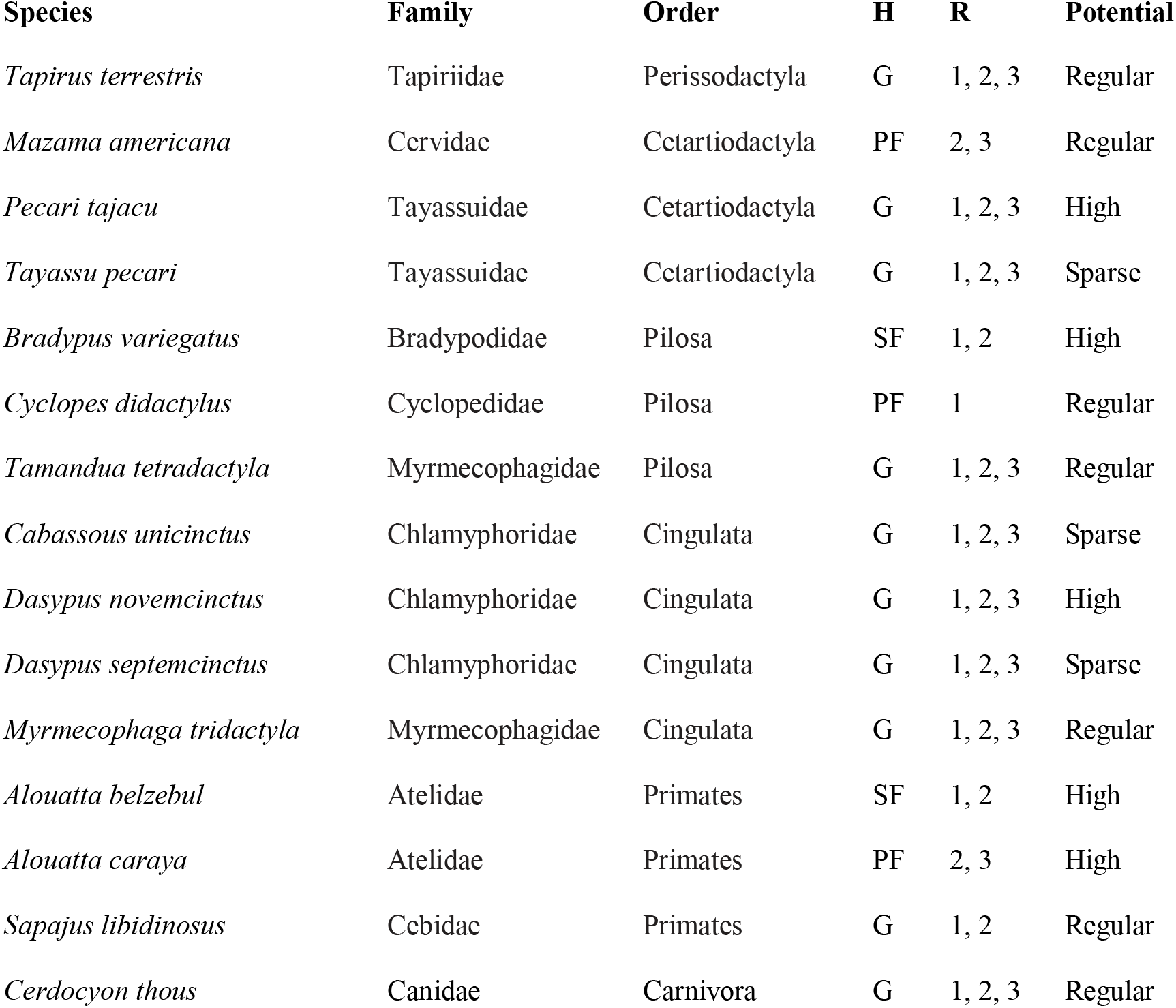
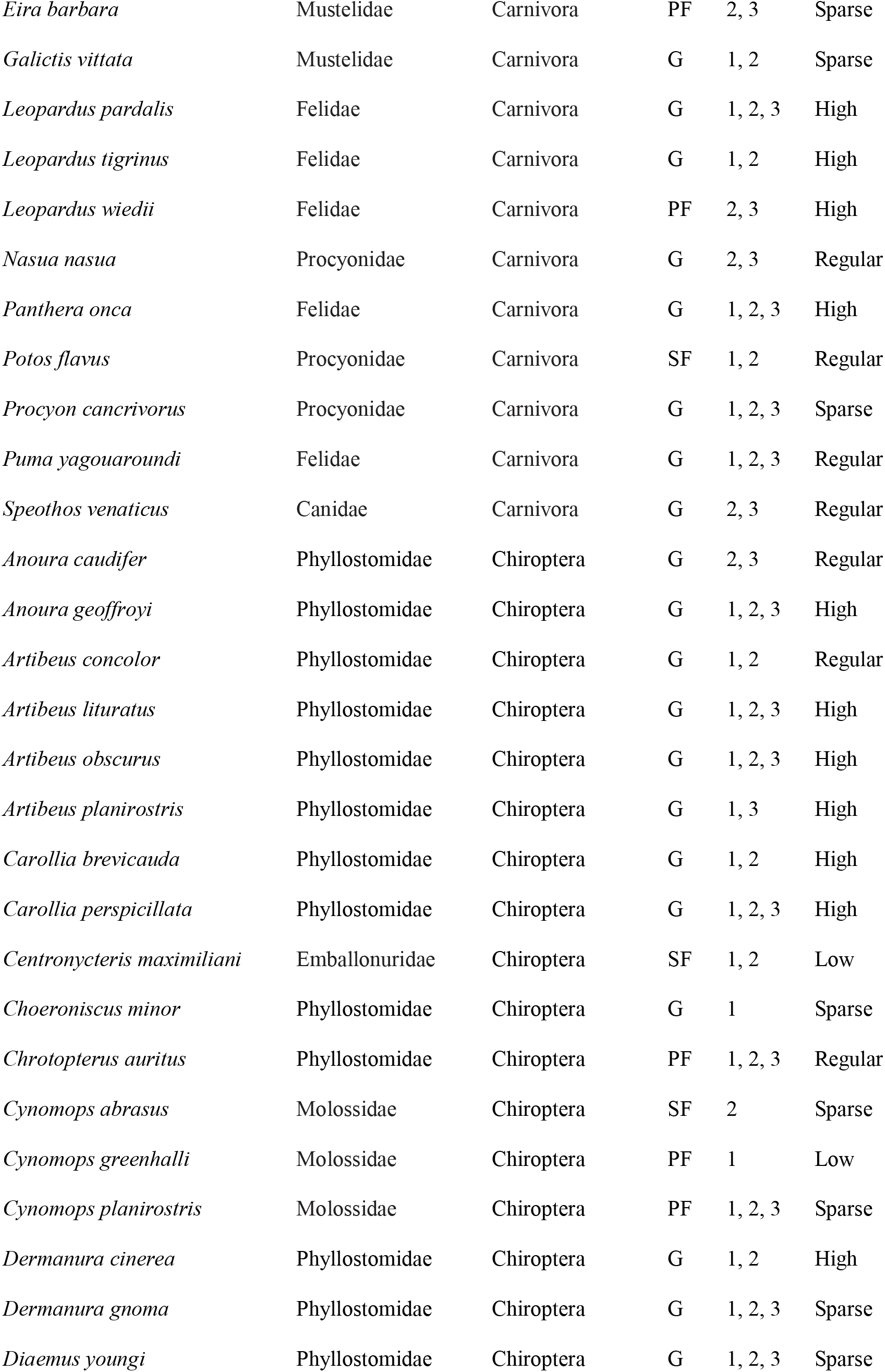
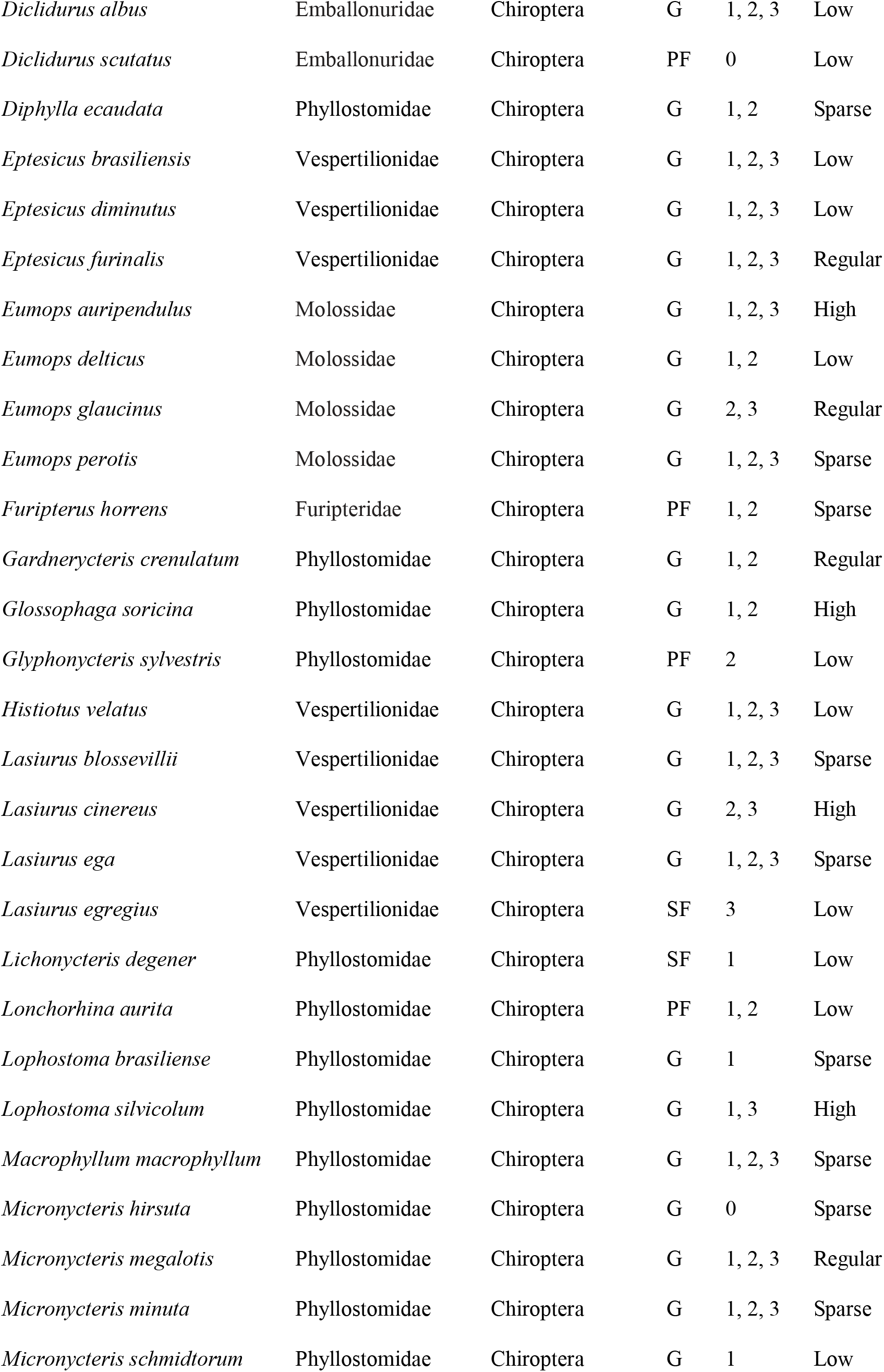
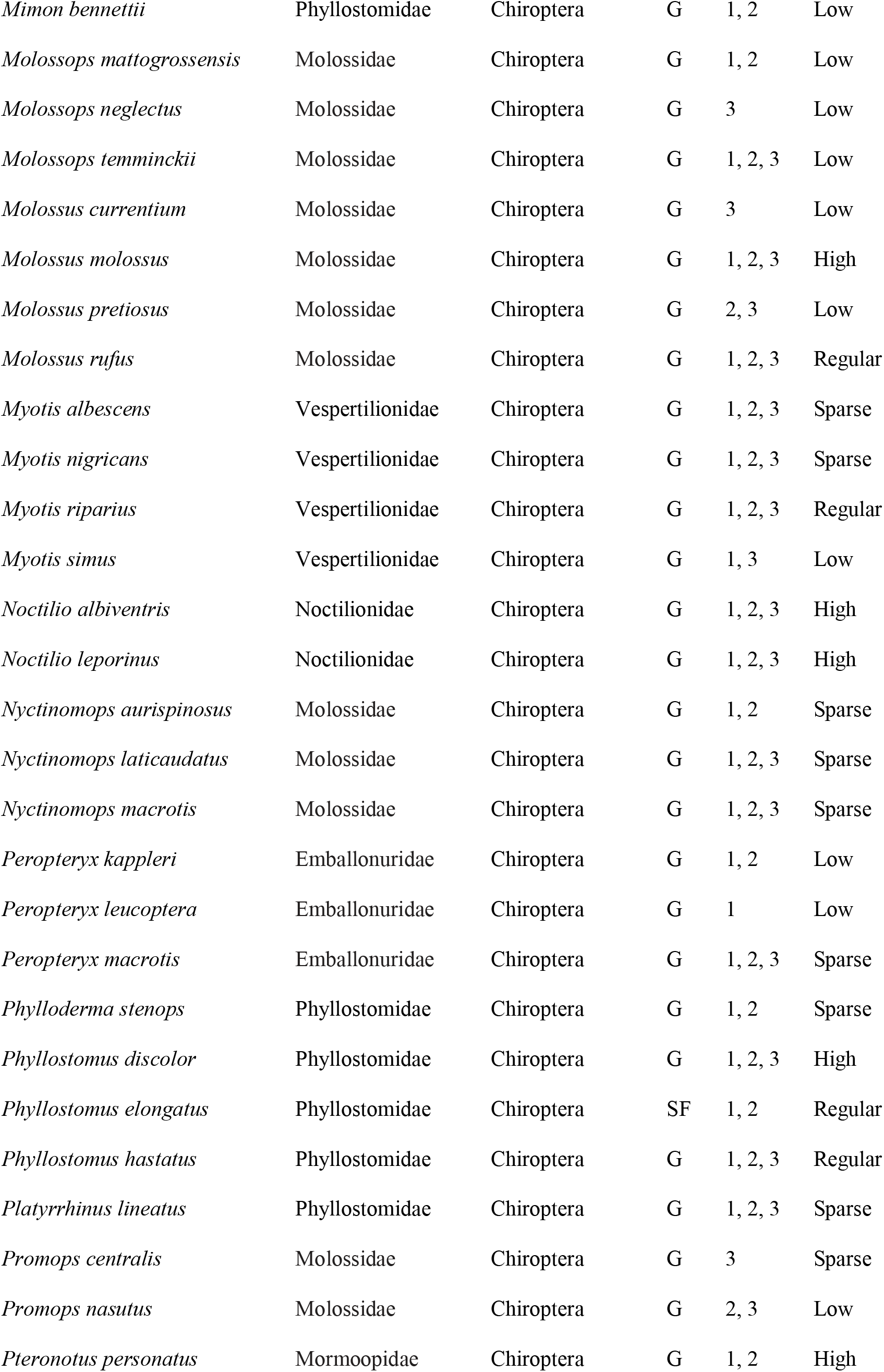
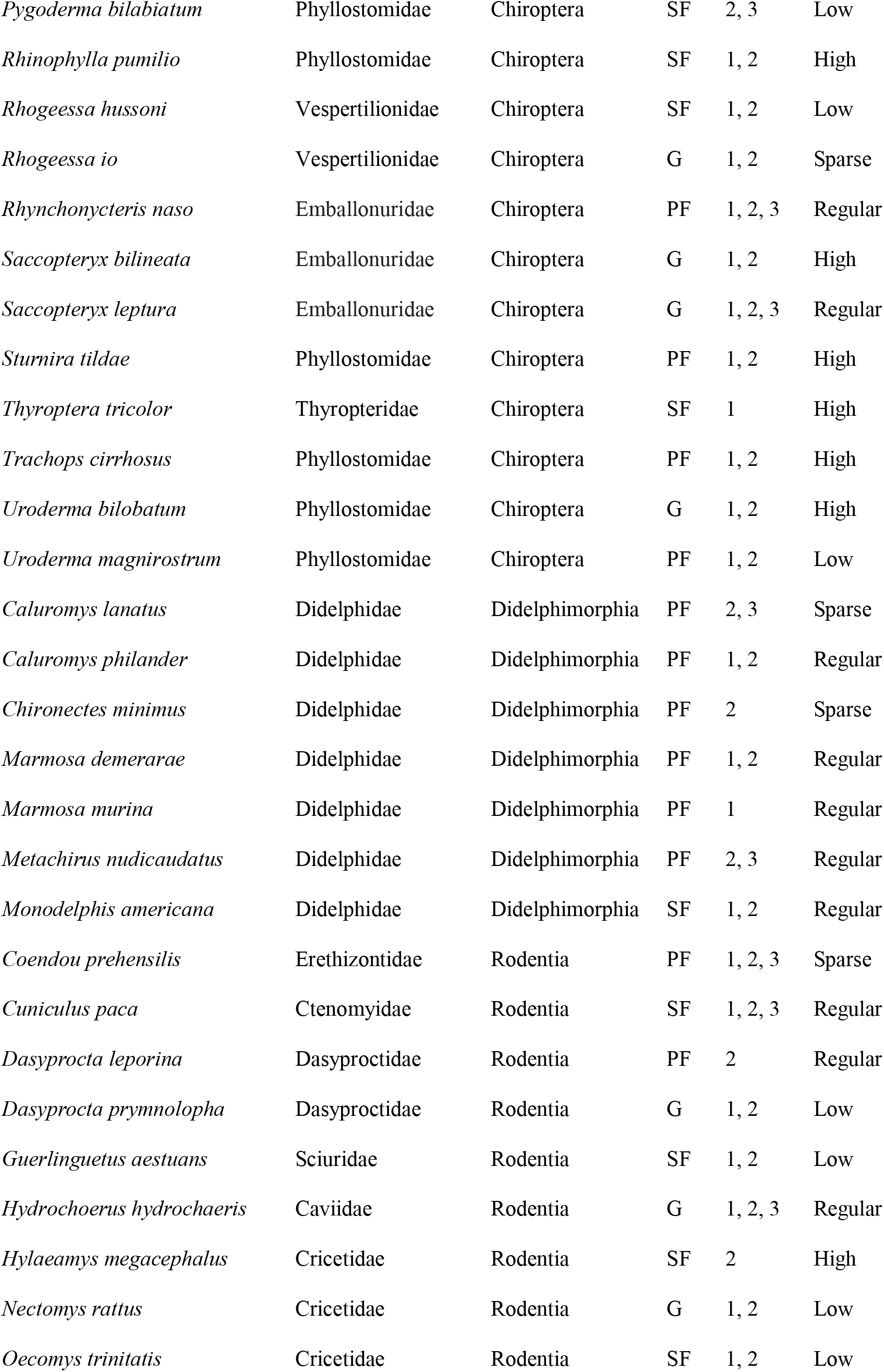
Potential mammal species for investigating the past connections between Amazonia and the Atlantic Forest. H = habitat preference (SF = Strict forest specialist; PF = Species that prefer forest; G = Generalists); R = Connective routes between Amazonia and the Atlantic Forest based on the IUCN expert species distribution maps (0 = Unidentified route; 1 = Northeast route [NE]; 2 = Central route [CE]; 3 = Southwest route [SW]); Potential = quantity of availability of genetic data based on the total number of nucleotide DNA sequences available on Genbank online database. The categories run from low (0-22 DNA sequences), to sparse (23-74 DNA sequences), satisfactory (75-225 DNA sequences), and high (226 to >1000 DNA sequences).

### Genetic data

We compiled genetic data for each species from the Genbank database [25]. Data compilation was done during January 2020, by registering the amount of molecular data available (nucleotide sequences) for each species. The genetic data was used to assess the taxonomic representativeness (i.e., which taxonomic groups represent the highest availability of published genetic data) and, consequently, their potential usefulness in evaluating the past existence and use of connections between Amazonia and the Atlantic Forest.

We created categories for the availability of genetic data to assess the potential usefulness of the mammal species. These categories were defined using the quantile function of the R package ‘stats’ [28]. We considered species with low availability of genetic data those with zero to 22 nucleotide sequences in the database: species with sparse genetic data availability those with 23 to 74 sequences, species with regular availability those with 75 to 225 sequences in the database and species with high data availability those with 226 to over 1000 sequences. We also calculated the average of sequences by species for each taxonomic order to compare the availability of genetic data for these different taxonomic groups.

### Identification of potential connective routes

To identify the potential connective routes between Amazonian and Atlantic forests used by mammals, the geographical areas of these connections on ecoregion maps were first delimited using Olson’s ecoregion polygons [27]. The NE route area was delimited using the boundaries of the Caatinga ecoregion, the transition areas between the Caatinga, the northern Cerrado and eastern Amazonia, the Babaçu Forest, and adjacent dry forests, which represent transition areas between Amazonia and the Atlantic Forest (Fig 1; route 1). The area of the CE route was selected using the limits of the Cerrado ecoregion (excluding the northern part, which was selected for the NE route; Fig 1; route 2). The area of the SW route was delimited using the boundaries of the Pantanal and the Chaco ecoregions, the transition areas between Amazonia and the Pantanal (such as Chiquitano Dry Forests), the southern Cerrado and the southwestern Atlantic Forest (Fig 1; route 3).

**Figure 1.**
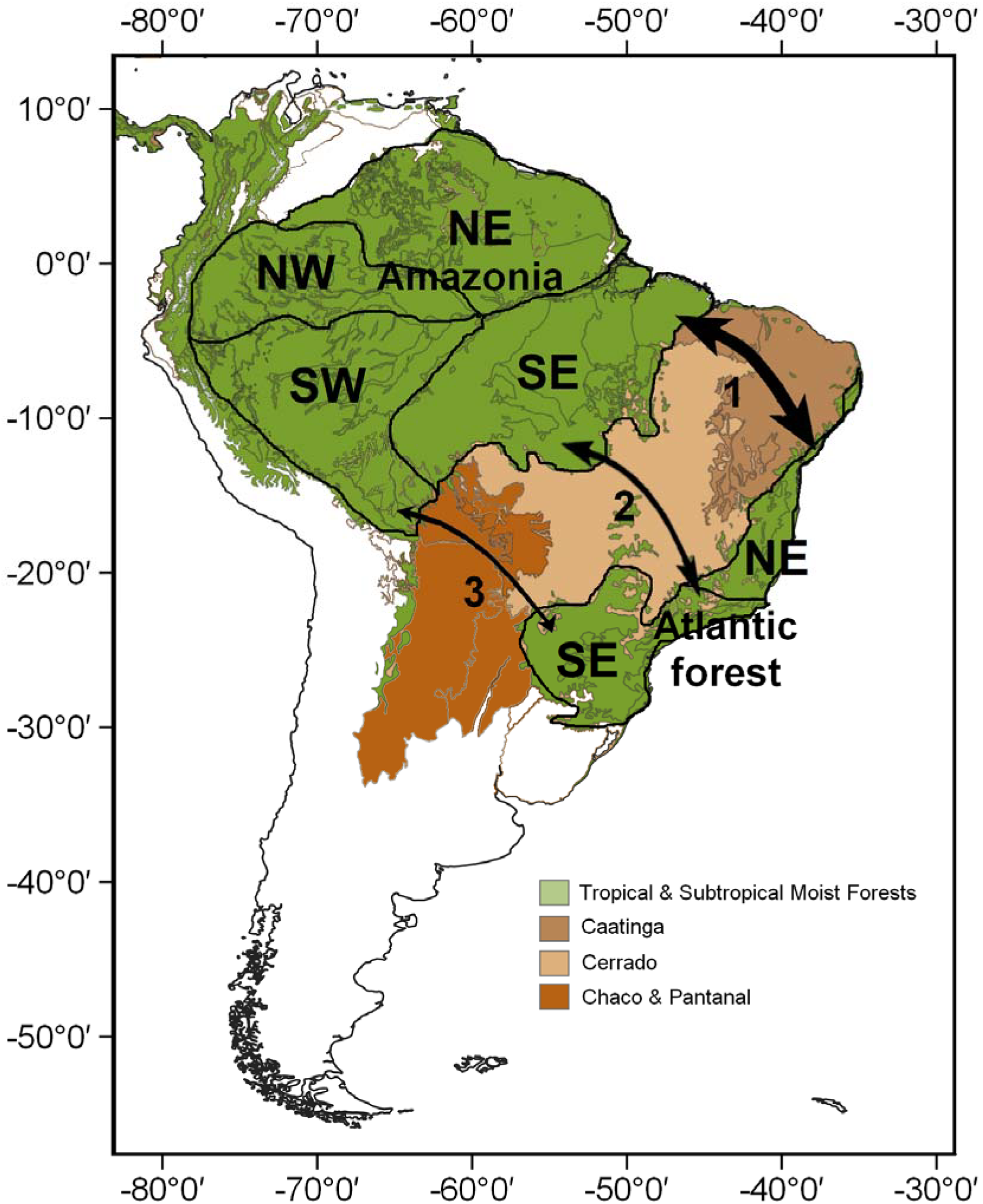
Distribution of Tropical Moist Forests in South America in green emphasizing Amazonia and the Atlantic Forest. Internal arrows represent connective routes between these forests through the Northeast/NE route (1), the Central/CE route (2) and the Southeast-Northwest/SW route (3). The width of the arrows represents the potential frequency that we found by which the routes have existed in the past based on distribution polygons of mammal species.

To explore how many species might have used each connective route, the delimited area for each route (NE, CE, and SW) was intersected with the species distribution maps using the function *gIntersection* in the R package ‘rgeos’ v. 0.5-5 [27]. Thereby, we quantified the total number of species associated with each route and the number of routes associated with each species. To visualize this result in the geographic space, we calculated the sum of rasters using the R package ‘raster’ v. 3.3 [39].

## Results

We compiled geographic distribution maps, information about habitat preferences, and genetic data for 127 mammal species occurring in both Amazonia and Atlantic Forest, following nine taxonomic orders: Didelphimorphia (7), Pilosa (3), Cingulata (4), Perissodactyla (1), Cetartiodactyla (3), Primates (3), Carnivora (12), Chiroptera (85) and Rodentia (9) (Table 1). According to the IUCN, the geographic distribution of 113 of those species appear to be continuous between Amazonia and the Atlantic Forest, whereas the remaining 14 species present disjunct distributions (S1 Table).

In terms of habitat use, 17 species were classified as strict forest specialists (SF), 23 species with preference for forests (PF) and 88 species as generalists (G) (Fig 2; Table 1). For the NE route, two species were SF, three PF and four were G. For the CE route, two species were SF, two PF and none were G. For the SW route, one species was SF, no one PF and two species were G (Fig 2; Table 1). For the NE and CE routes together, 10 species were SF, seven PF and 20 were G. For the three connective routes combined, one species was SF, four PF and 49 were G (Fig 2; Table 1). The result of the chi-square test for the habitat preference was significant (Chi-squared2 = 35.904, df = 12, p-value = 0.0003). Thus, we reject the null hypothesis, which states that habitat preference is independent from choice of connective route.

**Figure 2.**
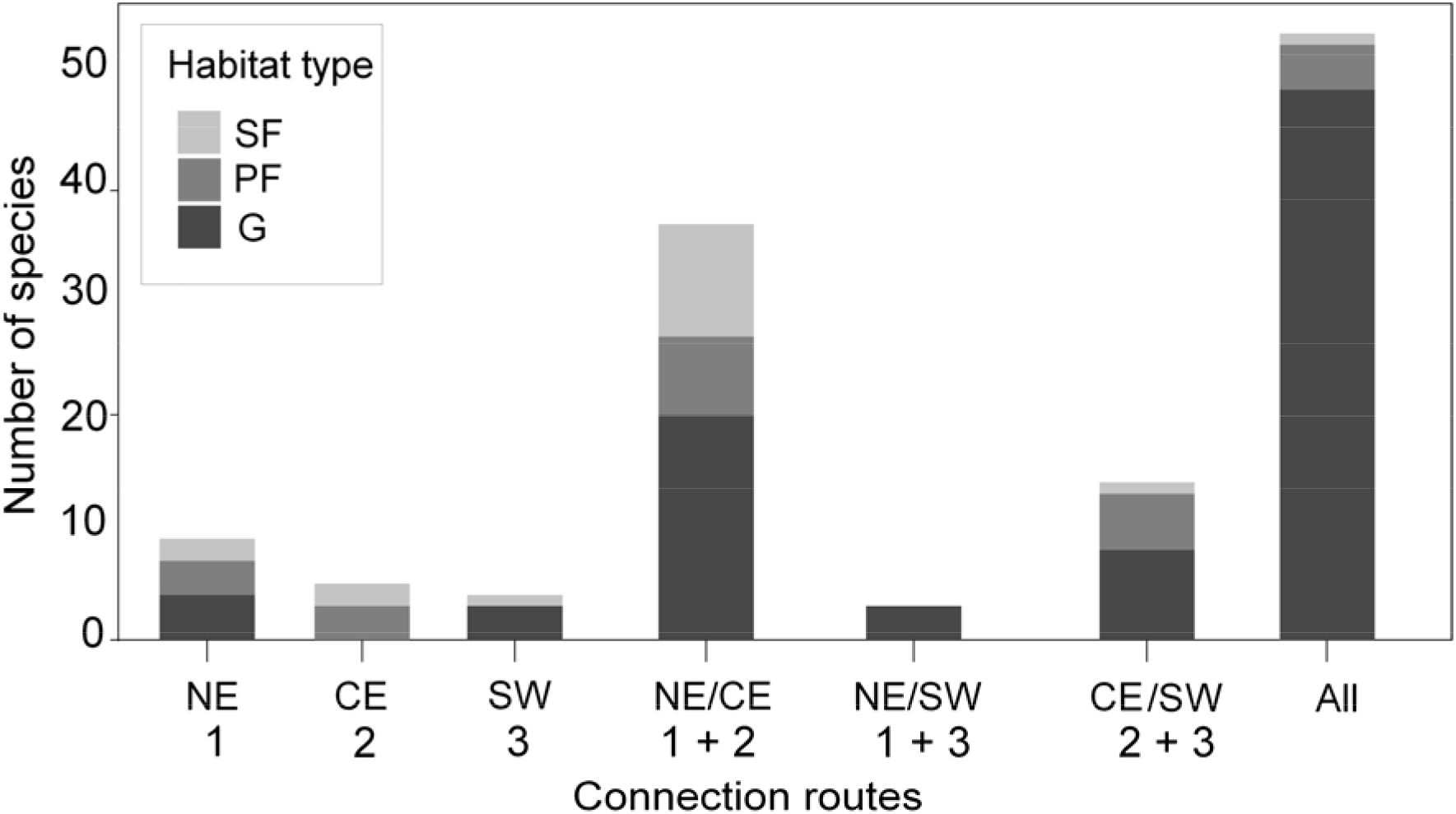
Number of mammalian species assumed to have dispersed by each of the connective routes between Amazonia and the Atlantic Forest, as evidenced by the species geographical distributions. The routes potentially used by mammalian species are presented along the x-axis: 1) NE = Northeast route, 2) CE = Central route, 3) SW = Southeast - Northwest route, and combinations of routes (1 + 2, 1 + 3, 2 + 3, and “All” for 1 + 2 + 3). The grayscale represents species habitat preferences where SF = Strict forest specialists, PF = Species that prefer forests and G = Generalists.

Most of the species identified in this study as potentially useful for assessing the connections between Amazonia and the Atlantic Forest have large amounts of genetic data available in the investigated database, including different molecular markers (Fig 3; Table 1; Genbank access link for details on available molecular data, in S1 Table). Thirty of the investigated species showed Low availability of genetic data in Genbank, 32 species show Sparse genetic data availability, 33 Regular and 32 High availability of genetic data (Fig 3; Table 1). The Cingulata and Chiroptera contained most of the available genetic data (with averaging about 1,000 and 590 sequences per species, respectively; Fig 4), followed by the Pilosa, Primates, Cetartiodactyla, Perissodactyla (averaging about 330, 300, 220, 200 and 190 sequences per species respectively; Fig 4). The Didelphimorphia and Rodentia also showed a considerable number of available nucleotide sequences (averaging about 115 and 93 sequences per species, respectively; Fig 4).

**Figure 3.**
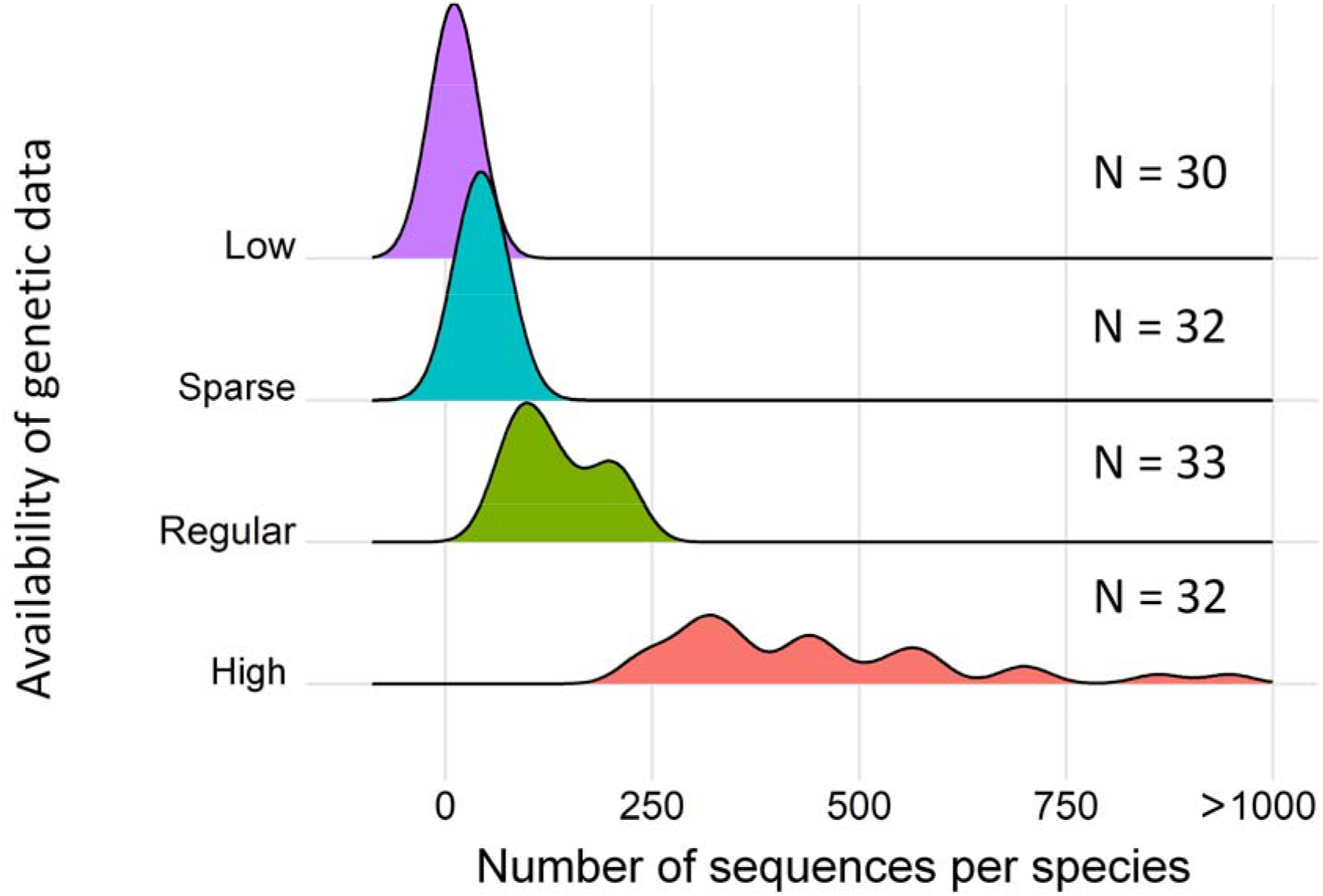
Availability of genetic data for mammalian species that occur in both Amazonia and the Atlantic Forest divided into the categories: Low = species with low availability of genetic data (from 0 to 22 sequences), Sparse = species with sparse availability of genetic data (from 23 to 74 sequences), Regular = species with satisfactory availability of genetic data (from 75 to 225 sequences), and High = species with high availability of genetic data (from 226 to over 1000 sequences).

**Figure 4.**
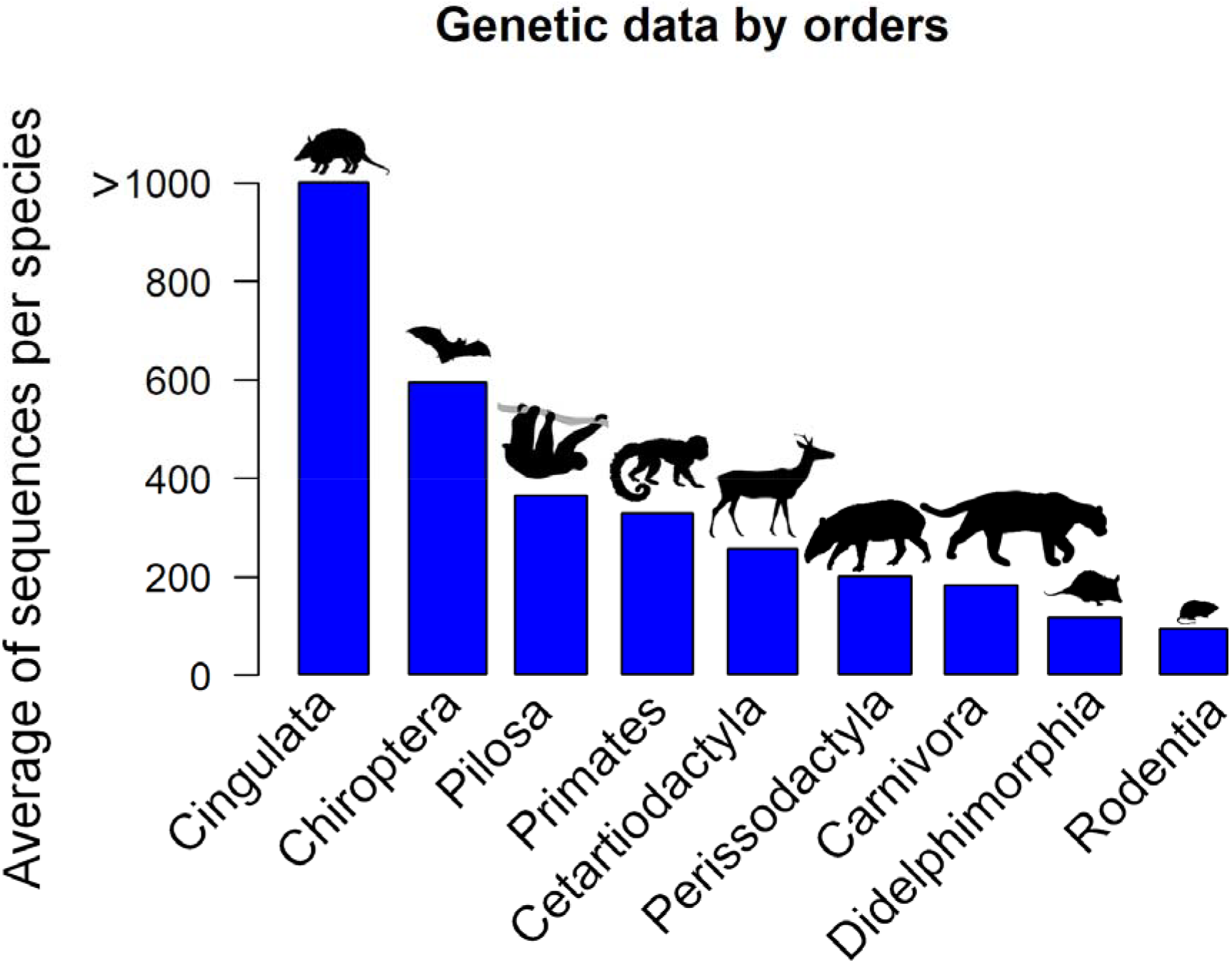
Availability of genetic data (Average of sequences per species) by orders for species of forest mammals that occur in both Amazonia and the Atlantic Forest.

As evidenced by the IUCN geographic distribution maps, 52 (41.6%) of the 127 evaluated species of mammals potentially dispersed between Amazonia and the Atlantic Forest using all three connective routes (NE, CE, and SW). Thirty-eight of the 127 species (30.4%) potentially used both the NE and the CE routes, whereas 14 species (11.2%) may have potentially used the CE and the SW routes. Nine species (7.2%) are expected to have used the NE route exclusively, five species (4%) the CE route exclusively, and four species (3.2%) the SW route exclusively. Only three species show potential to have used a combination of the NE and the SW routes (2.4%) (Fig 5). Hence, the potentially most used connections were the NE route followed by the CE route (Fig 1, Fig 5, Fig 6). For two bat species *Diclidurus scutatus* and *Micronycteris hirsute* it was not possible to attribute a potential connective route, as these present extremely disjunct distributions between Amazonia and Atlantic Forest (Table 1).

**Figure 5.**
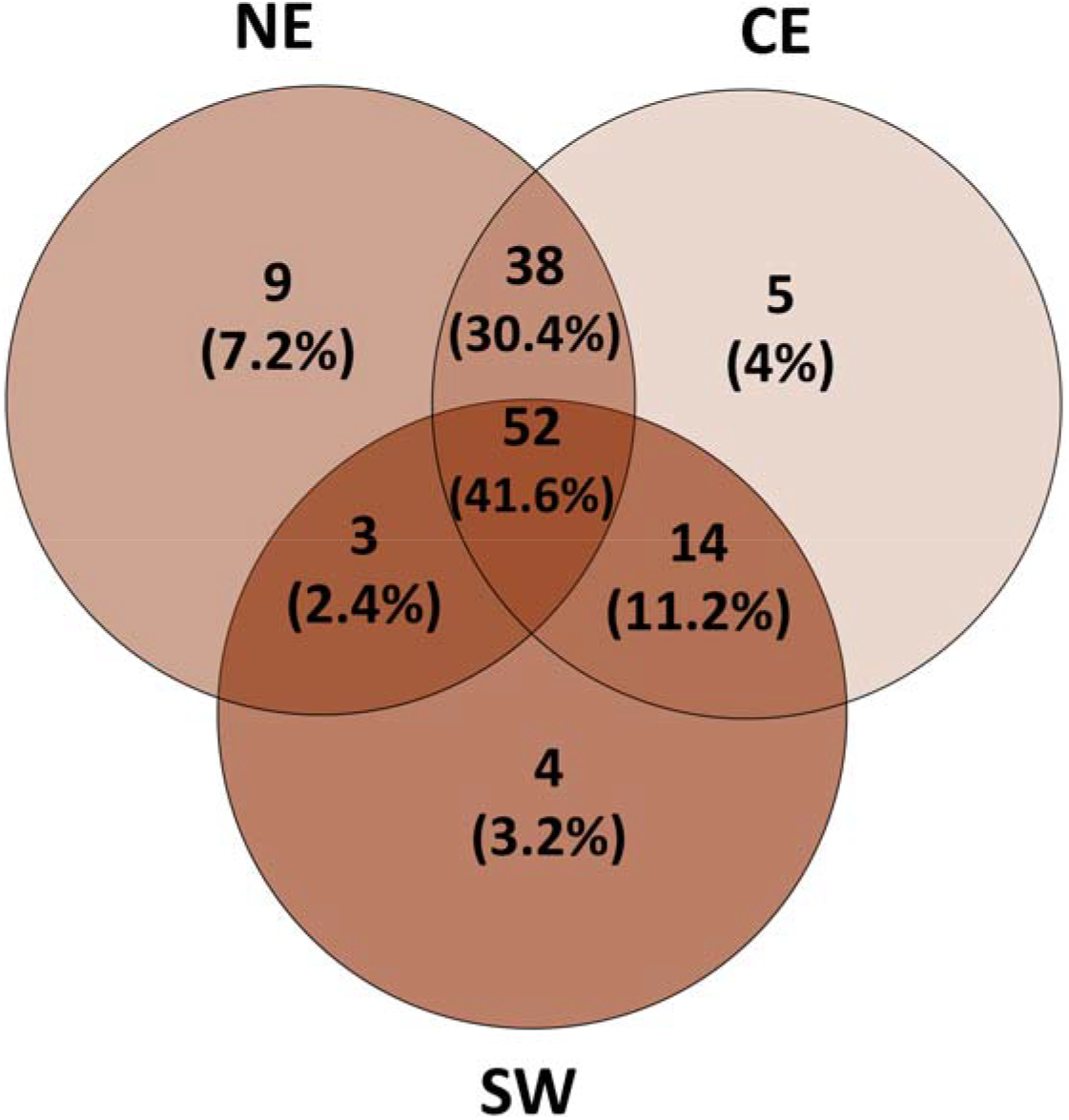
Venn diagram showing the past connective routes between Amazonia and the Atlantic Forest as evidenced by the IUCN geographical distribution maps for mammalian species that occur in both biomes.

**Figure 6.**
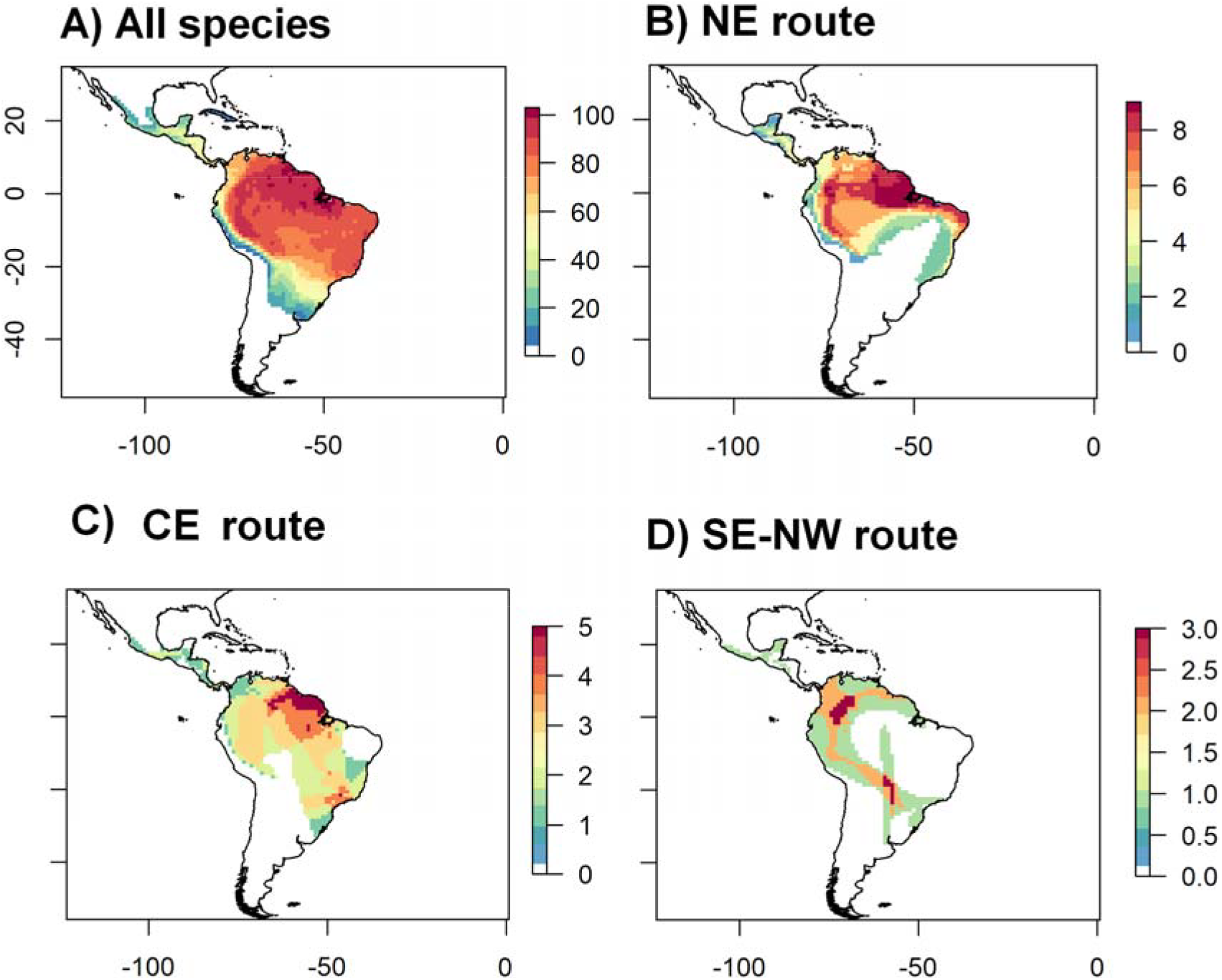
The overlaid distribution maps of potential mammal species for investigating the connections between Amazonia and the Atlantic Forest show most connections for the Northeast route (NE route), followed by the Central route (CE route) and finally the smallest number of species were associated with the Southeast - Northwest route (SW route). The colour scale represents the number of species per pixel on the map. A) Overlap in the species distribution maps for all mammal species sampled in this study; B) Overlap in species distribution maps for species associated with the NE route; C) Overlap in species distribution maps for species associated with the CE route; D) Overlap in species distribution maps for species associated with the SW route.

## Discussion

Here we present an unprecedented list of potential mammal species for investigating the past connections between Amazonia and the Atlantic Forest. Previous studies, including data compilations, analysed a limited number of species of mammals. Here we were able to include 127 species of mammals occurring in in both Amazonia and the Atlantic Forest to add information about the potential past connections between the two biomes. We compiled information about the species’ distributions, habitat preferences, and the quantity of genetic data available. We also showed the possible routes used by these mammals, and the association between habitat use and their potential connection route between the two rainforest regions.

According to Por [5], the SW connection between Amazonia and the Atlantic Forest was the most used route. The connection through the NE was the second most common route, whereas the route through the riverine forests in the interior of the Cerrado (CE route) was the least used. This hypothesis has been confirmed by a review of vertebrate molecular data [10]. The single previous study compiling data for multiple species of mammals evaluating these connective routes was Costa [6]. This author investigated the phylogeography of small mammals and found that the CE route was potentially the most used route. However, our results show that, differently from these previous studies, the NE route would have been the most common route. The CE route was the second most common and the SW route was the least used. These contrasting results highlight the need for further studies including multiple taxa, ecological traits, and evolutionary data. The list we compiled here has the potential to subsidize other mammal phylogeographic studies and shed light on the temporal and spatial use of the connections between Amazonia and the Atlantic forest in relation to South American mammals’ ecology and evolution.

In fact, studies on paleo vegetation, pollen data and biogeographic approaches have shown evidence of past connections between the northern Atlantic Forest and the eastern Amazonia through the NE route [7, 12–14, 20, 40–42). Even so, no had directly shown the potential importance of this route in connecting Amazonia and the Atlantic Forest. Also, several species showed evidence of having used more than one route in the past. Most species potentially used both the NE and CE routes or even the whole of the three routes.

Previous studies underestimated the number of mammalian species that could have dispersed between Amazonia and the Atlantic Forest, especially for the NE route, but even our study is likely to underestimate this number. For example, sampling artefacts deficiencies in the dry diagonal of the northeast, in addition to the non-inclusion of point records published for the northeast or available from specimens deposited in mammal collections [16] may have led us to underestimate the number of potential taxa that could have dispersed through the NE route. Additionally, many species showing disjunct distributions, probably represent either sampling deficiencies, such as the two bat species *Diclidurus scutatus* and *Micronycteris hirsute*, and also the huge extension of range of *Promops centralis* after using complementary sampling schemes or cryptic diversity [43].

Examples of mammal species need to be reassessed regarding their past occurrences along the NE route, include the anteater *Cyclopes didactylus*, the Kinkajou *Potos flavus* and the squirrel *Guerlinguetus brasiliensis* [44–46, respectively]. The taxonomic status of the guariba *Alouatta ululata*, previously a subspecies of *A. belzebul*, has been elevated to the species category [47] and requires a revaluation through molecular and cytogenetic approaches, as suggested by Viana et al. [48]. Miranda et al. [49–50] presented new records of common forest species for the NE route, such as the marsupials *Marmosa demerarae*, *Metachirus nudicaudatus* and *Didelphis marsupialis* (personal obs. C.L. Miranda). The rodent *Oecomys catherinae*, considered restricted to the Atlantic Forest, has been recorded along the NE route [51, 52]. The black-eared opossum *D. marsupialis* presents disjunct distribution, but recent records point to no disjunction along the NE route [e.g., 49, 50, 53]. The three-striped short-tailed opossum, *Monodelphis americana*, is also present in the eastern Amazonia and in the northeastern Atlantic Forest, with the populations south of the São Francisco River belonging to another species of the genus *Monodelphis* [54]. Hence, updating the known distribution of these species and others would certainly increase the number of species showing evidence of the past connections between Amazonia and the Atlantic Forests, mainly through the NE route. Additionally, as new records appear with increased sampling and systematic biogeographic studies from the NE region, future reassessments will certainly find further species showing evidence of past dispersions along the NE route [e.g., 47, 48, 53].

As highlighted by Costa [6], the forest environments in the Cerrado ecoregion should function as ecological [55] and historical corridors, allowing forest species to be present in the region. Thus, these forests form a connection that until our study had not been specifically recognized; here we call it the CE route, although it had already been considered a potentially independent route in other studies [5, 6, 15, 17]. Besides those riparian forests in the northeast dry diagonal, along the NE route, there is a marked presence of other types of forested habitats that may also serve the same function. Some examples include the Babaçu forests, the semi-deciduous forests, the mangroves, and the “boqueirões” (humid forests that occur in patches in the Caatinga, which some authors consider remnants of the Atlantic Forest, while others consider relic environments with biogeographical composition and affinities yet to be properly understood) and the “brejos de altitude” or Altitude Swamps [49]. To evaluate this hypothesis would, however, require specific studies that focus on the species that show evidence of the NE route to better map their past dispersion routes.

According to the IUCN species distribution maps, many mammalian species seem to have continuous distributions between Amazonia and the Atlantic Forest through deciduous and semi-deciduous forests in the interior of the dry diagonal (composed by the Caatinga, the Cerrado and the Chaco). However, many phylogeographic studies reveal that some of these species, with seemingly continuous distributions, consist of currently isolated populations [e.g., 56]. Moreover, many isolated populations show evolutionary well-structured lineages with significant genetic divergence, so their taxonomic status is probably in need of revision as they could represent species complexes [e.g., 6, 34, 56, 58]. Hence, further phylogeographic studies are necessary if we are to reveal whether these mammalian populations are indeed connected or isolated in function of the current fragmentation pattern of the South American forests, particularly that of anthropic origin so evident in the Atlantic Forest remnants and in the arc of deforestation of Amazonia. Indeed, the landscape becomes increasingly fragmented as we continue to lose forest habitat, the environmental protection system fails to connect the ecoregions between Amazonia and the Atlantic Forest [59], and under the complete laxation of the Brazilian environmental laws under the current federal government [60]. Given the relationship that we observed between species’ habitat preferences and their potential connection routes in the past, such forest loss and habitat fragmentation could have dire consequences for the mammal populations along these dispersal pathways.

Whereas the CE route was associated with forest specialists or species mostly with forest habits, most routes between Amazonia and the Atlantic Forest were associated with habitat generalists. The NE route, for example, passed through the Caatinga or the Cerrado/Caatinga transition, which is a region of high environmental heterogeneity [61, 62]. Thus, here, we found species with a greater variety of habitat preferences. Subsequently, we believe that the inclusion of species with a wider range of habitat preferences (i.e. not only forest specialists) could be decisive for the identification and quantification of the past use of connective routes between Amazonia and the Atlantic Forest; not least because the past use of the connective routes seems related to the environmental heterogeneity of each region and environmental changes in the past [11, 12, 13]. Therefore, it would be interesting to make use of the 127 species that we identified, to evaluate the past forest connections in relation to different sampling methods and the differences in ecological characteristics among taxonomic groups.

The availability of genetic data for our 127 candidate species, revealed that many of them would serve for assessing the existence and importance of the past connections between Amazonia and the Atlantic Forest. Also, although we only show the total amount of available nucleotide sequences for each species, this initial compilation can be extremely useful in facilitating more detailed evaluations for future phylogeographic simulations and to further explore the past connections between Amazonia and the Atlantic Forest. However, it is important to highlight that, although many of these species have a large amount of available genetic data, it is common for many of the nucleotide sequences to contain missing information, which could render them unfit for use. Even so, many recent studies provide complete genetic data for mammals favouring the development of phylogeographic studies 57, 58, 63, 64]. Therefore, our results, not only inform us about the dispersal history of the 127 mammalian species among the humid Neotropical forests, but our results also bring new insights about the connective routes that may have once existed between Amazonia and the Atlantic Forest.

## Acknowledgments

We thank Renan Maestri, Fabricio Villalobos, Thales R. O. de Freitas and Yennie K. Bredin for their critical revision of this manuscript. AFM was supported by a doctoral fellowship (grant 141008/2016-4) provided by the Conselho Nacional de Desenvolvimento Científico e Tecnológico (CNPq). LD is a member of the National Institute for Science and Technology (INCT) in Ecology, Evolution and Biodiversity Conservation, supported by MCTIC/CNPq (proc. 465610/2014-5) and FAPEG (proc. 201810267000023)., LD and MJRP were supported by CNPq Productivity Fellowships (grants 307527/2018-2 and 311297/2018-8). CDR thanks the financial support from Alexander von Humboldt Foundation.

## Authors contribution

AFM, LD and MJRP designed the study. AFM compiled the data and CLM and MJRP revised and complement it. AFM and CDR analyses the data. AFM, wrote the manuscript with contribution of LD, CDR, CLM and MJRP.

## Availability of data material

Not applicable.

